## Supplementary Information for "StressME: unified computing framework of *Escherichia coli* metabolism, gene expression, and stress responses"

### Supplemental information

#### SI-1: Difference between StressME and EcoliME [9]

##### Table S1 Difference in model components between StressME and EcoliME

| Model components | StressME | EcoliME |
|---|---|---|
| **genes** | 1,689 | 1,678 |
| **mRNAs** | 1,689 | 1,678 |
| **tRNAs** | 63 | 63 |
| **proteins** | 1,578 | 1,568 |
| **protein folding states** | 9,561 | 0 |
| **metabolites** | 1,673 | 1,671 |
| **complexes** | 1,692 | 1,526 |
| **reactions** | 36,735 | 12,655 |

#### SI-2: Temperature-dependent thermostability, aggregation propensity, and folding rate constants for ten folding proteins added to StressME.

##### Table S2 Temperature-dependent thermostability (Oobatake Keq [27]) for 10 folding proteins added to StressME

| | b0605 | b0606 | b2962 | b4209 | b3662 | b0812 | b3961 | b4062 | b4063 | b0439 |
|---|---|---|---|---|---|---|---|---|---|---|
| **26℃** | 5.0E-12 | 4.2E+03 | 7.1E-07 | 8.6E+00 | 1.1E-29 | 8.4E+02 | 2.1E-04 | 2.1E-02 | 1.4E+07 | 9.5E+31 |
| **27℃** | 8.0E-12 | 6.2E+03 | 8.4E-07 | 1.1E+01 | 4.4E-29 | 8.9E+02 | 3.0E-04 | 2.1E-02 | 1.2E+07 | 8.1E+31 |
| **28℃** | 1.3E-11 | 9.6E+03 | 1.0E-06 | 1.4E+01 | 1.8E-28 | 9.7E+02 | 4.4E-04 | 2.2E-02 | 1.1E+07 | 7.5E+31 |
| **29℃** | 2.2E-11 | 1.6E+04 | 1.2E-06 | 1.8E+01 | 7.4E-28 | 1.1E+03 | 6.7E-04 | 2.3E-02 | 9.7E+06 | 7.6E+31 |
| **30℃** | 3.8E-11 | 2.7E+04 | 1.5E-06 | 2.4E+01 | 3.2E-27 | 1.2E+03 | 1.1E-03 | 2.4E-02 | 8.8E+06 | 8.4E+31 |
| **31℃** | 6.6E-11 | 4.8E+04 | 1.8E-06 | 3.2E+01 | 1.5E-26 | 1.4E+03 | 1.7E-03 | 2.6E-02 | 8.1E+06 | 1.0E+32 |
| **32℃** | 1.2E-10 | 9.1E+04 | 2.2E-06 | 4.5E+01 | 6.9E-26 | 1.6E+03 | 2.9E-03 | 2.8E-02 | 7.5E+06 | 1.3E+32 |
| **33℃** | 2.1E-10 | 1.8E+05 | 2.7E-06 | 6.3E+01 | 3.4E-25 | 1.9E+03 | 5.0E-03 | 3.0E-02 | 7.2E+06 | 1.8E+32 |
| **34℃** | 3.7E-10 | 3.7E+05 | 3.4E-06 | 9.1E+01 | 1.7E-24 | 2.3E+03 | 8.9E-03 | 3.4E-02 | 6.9E+06 | 2.8E+32 |

| | | | | | | | | | | |
|---|---|---|---|---|---|---|---|---|---|---|
| **35℃** | 6.9E-10 | 8.1E+05 | 4.3E-06 | 1.3E+02 | 9.1E-24 | 2.8E+03 | 1.6E-02 | 3.8E-02 | 6.8E+06 | 4.5E+32 |
| **36℃** | 1.3E-09 | 1.8E+06 | 5.4E-06 | 2.0E+02 | 5.0E-23 | 3.4E+03 | 3.1E-02 | 4.3E-02 | 6.7E+06 | 8.0E+32 |
| **37℃** | 2.4E-09 | 4.4E+06 | 6.9E-06 | 3.1E+02 | 2.8E-22 | 4.3E+03 | 5.9E-02 | 4.9E-02 | 6.8E+06 | 1.5E+33 |
| **38℃** | 4.7E-09 | 1.1E+07 | 8.9E-06 | 4.9E+02 | 1.7E-21 | 5.5E+03 | 1.2E-01 | 5.7E-02 | 7.0E+06 | 3.1E+33 |
| **39℃** | 9.1E-09 | 2.9E+07 | 1.2E-05 | 7.8E+02 | 1.0E-20 | 7.2E+03 | 2.4E-01 | 6.6E-02 | 7.2E+06 | 6.8E+33 |
| **40℃** | 1.8E-08 | 7.9E+07 | 1.5E-05 | 1.3E+03 | 6.3E-20 | 9.5E+03 | 5.1E-01 | 7.8E-02 | 7.6E+06 | 1.6E+34 |
| **41℃** | 3.6E-08 | 2.2E+08 | 2.0E-05 | 2.1E+03 | 4.1E-19 | 1.3E+04 | 1.1E+00 | 9.4E-02 | 8.2E+06 | 4.1E+34 |
| **42℃** | 7.4E-08 | 6.7E+08 | 2.6E-05 | 3.5E+03 | 2.8E-18 | 1.7E+04 | 2.4E+00 | 1.1E-01 | 8.9E+06 | 1.1E+35 |
| **43℃** | 1.5E-07 | 2.1E+09 | 3.5E-05 | 6.1E+03 | 1.9E-17 | 2.4E+04 | 5.6E+00 | 1.4E-01 | 9.8E+06 | 3.3E+35 |
| **44℃** | 3.2E-07 | 6.7E+09 | 4.7E-05 | 1.1E+04 | 1.4E-16 | 3.3E+04 | 1.3E+01 | 1.7E-01 | 1.1E+07 | 1.0E+36 |
| **45℃** | 6.7E-07 | 2.3E+10 | 6.3E-05 | 1.9E+04 | 1.0E-15 | 4.7E+04 | 3.1E+01 | 2.1E-01 | 1.2E+07 | 3.5E+36 |
| **46℃** | 1.4E-06 | 7.9E+10 | 8.6E-05 | 3.4E+04 | 8.0E-15 | 6.7E+04 | 7.7E+01 | 2.6E-01 | 1.4E+07 | 1.2E+37 |

#### Table S2 Aggregation propensity for 10 folding proteins added to StressME

| | propensity | confidence |
|---|---|---|
| **b0605** | 5 | 5.29 |
| **b0606** | 12 | 5.19 |
| **b2962** | 2 | 7.31 |
| **b4209** | 2 | 5.54 |
| **b3662** | 14 | 5.60 |
| **b0812** | 6 | 5.65 |
| **b3961** | 9 | 5.18 |
| **b4062** | 3 | 5.14 |
| **b4063** | 3 | 6.21 |
| **b0439** | 11 | 5.12 |

#### Table S3 Temperature-dependent folding rate constants for ten folding proteins added to StressME

| | b0605 | b0606 | b2962 | b4209 | b3662 | b0812 | b3961 | b4062 | b4063 | b0439 |
|---|---|---|---|---|---|---|---|---|---|---|
| **26℃** | 3.2E-02 | 1.4E+00 | 1.6E-02 | 1.8E+04 | 8.1E-03 | 2.5E+03 | 2.5E+04 | 1.9E+03 | 6.1E+04 | 1.9E+01 |
| **27℃** | 4.1E-02 | 1.7E+00 | 2.1E-02 | 2.3E+04 | 1.0E-02 | 3.2E+03 | 3.2E+04 | 2.4E+03 | 7.7E+04 | 2.4E+01 |
| **28℃** | 5.2E-02 | 2.2E+00 | 2.7E-02 | 2.9E+04 | 1.3E-02 | 4.0E+03 | 4.1E+04 | 3.1E+03 | 9.9E+04 | 3.1E+01 |
| **29℃** | 6.6E-02 | 2.8E+00 | 3.4E-02 | 3.7E+04 | 1.7E-02 | 5.2E+03 | 5.2E+04 | 3.9E+03 | 1.3E+05 | 3.9E+01 |
| **30℃** | 8.4E-02 | 3.6E+00 | 4.3E-02 | 4.7E+04 | 2.1E-02 | 6.6E+03 | 6.6E+04 | 5.0E+03 | 1.6E+05 | 5.0E+01 |
| **31℃** | 1.1E-01 | 4.6E+00 | 5.5E-02 | 6.0E+04 | 2.7E-02 | 8.3E+03 | 8.3E+04 | 6.4E+03 | 2.0E+05 | 6.3E+01 |
| **32℃** | 1.3E-01 | 5.8E+00 | 7.0E-02 | 7.6E+04 | 3.4E-02 | 1.1E+04 | 1.1E+05 | 8.1E+03 | 2.6E+05 | 8.0E+01 |
| **33℃** | 1.7E-01 | 7.3E+00 | 8.8E-02 | 9.7E+04 | 4.3E-02 | 1.3E+04 | 1.3E+05 | 1.0E+04 | 3.3E+05 | 1.0E+02 |
| **34℃** | 2.2E-01 | 9.3E+00 | 1.1E-01 | 1.2E+05 | 5.5E-02 | 1.7E+04 | 1.7E+05 | 1.3E+04 | 4.1E+05 | 1.3E+02 |
| **35℃** | 2.7E-01 | 1.2E+01 | 1.4E-01 | 1.5E+05 | 6.9E-02 | 2.1E+04 | 2.1E+05 | 1.6E+04 | 5.2E+05 | 1.6E+02 |
| **36℃** | 3.4E-01 | 1.5E+01 | 1.8E-01 | 1.9E+05 | 8.7E-02 | 2.7E+04 | 2.7E+05 | 2.1E+04 | 6.5E+05 | 2.0E+02 |
| **37℃** | 4.3E-01 | 1.9E+01 | 2.2E-01 | 2.4E+05 | 1.1E-01 | 3.4E+04 | 3.4E+05 | 2.6E+04 | 8.2E+05 | 2.5E+02 |
| **38℃** | 5.4E-01 | 2.3E+01 | 2.8E-01 | 3.1E+05 | 1.4E-01 | 4.2E+04 | 4.2E+05 | 3.2E+04 | 1.0E+06 | 3.2E+02 |
| **39℃** | 6.8E-01 | 2.9E+01 | 3.5E-01 | 3.8E+05 | 1.7E-01 | 5.3E+04 | 5.3E+05 | 4.1E+04 | 1.3E+06 | 4.0E+02 |

| | | | | | | | | | | |
|---|---|---|---|---|---|---|---|---|---|---|
| **40℃** | 8.5E-01 | 3.7E+01 | 4.4E-01 | 4.8E+05 | 2.2E-01 | 6.6E+04 | 6.7E+05 | 5.1E+04 | 1.6E+06 | 5.0E+02 |
| **41℃** | 1.1E+00 | 4.6E+01 | 5.5E-01 | 6.0E+05 | 2.7E-01 | 8.3E+04 | 8.3E+05 | 6.4E+04 | 2.0E+06 | 6.3E+02 |
| **42℃** | 1.3E+00 | 5.7E+01 | 6.9E-01 | 7.5E+05 | 3.4E-01 | 1.0E+05 | 1.0E+06 | 8.0E+04 | 2.5E+06 | 7.9E+02 |
| **43℃** | 1.7E+00 | 7.1E+01 | 8.6E-01 | 9.4E+05 | 4.2E-01 | 1.3E+05 | 1.3E+06 | 9.9E+04 | 3.2E+06 | 9.8E+02 |
| **44℃** | 2.1E+00 | 8.9E+01 | 1.1E+00 | 1.2E+06 | 5.2E-01 | 1.6E+05 | 1.6E+06 | 1.2E+05 | 3.9E+06 | 1.2E+03 |
| **45℃** | 2.6E+00 | 1.1E+02 | 1.3E+00 | 1.5E+06 | 6.5E-01 | 2.0E+05 | 2.0E+06 | 1.5E+05 | 4.9E+06 | 1.5E+03 |
| **46℃** | 3.2E+00 | 1.4E+02 | 1.7E+00 | 1.8E+06 | 8.1E-01 | 2.5E+05 | 2.5E+06 | 1.9E+05 | 6.1E+06 | 1.9E+03 |

### SI-3: Temperature-dependent phenotypes and proteome

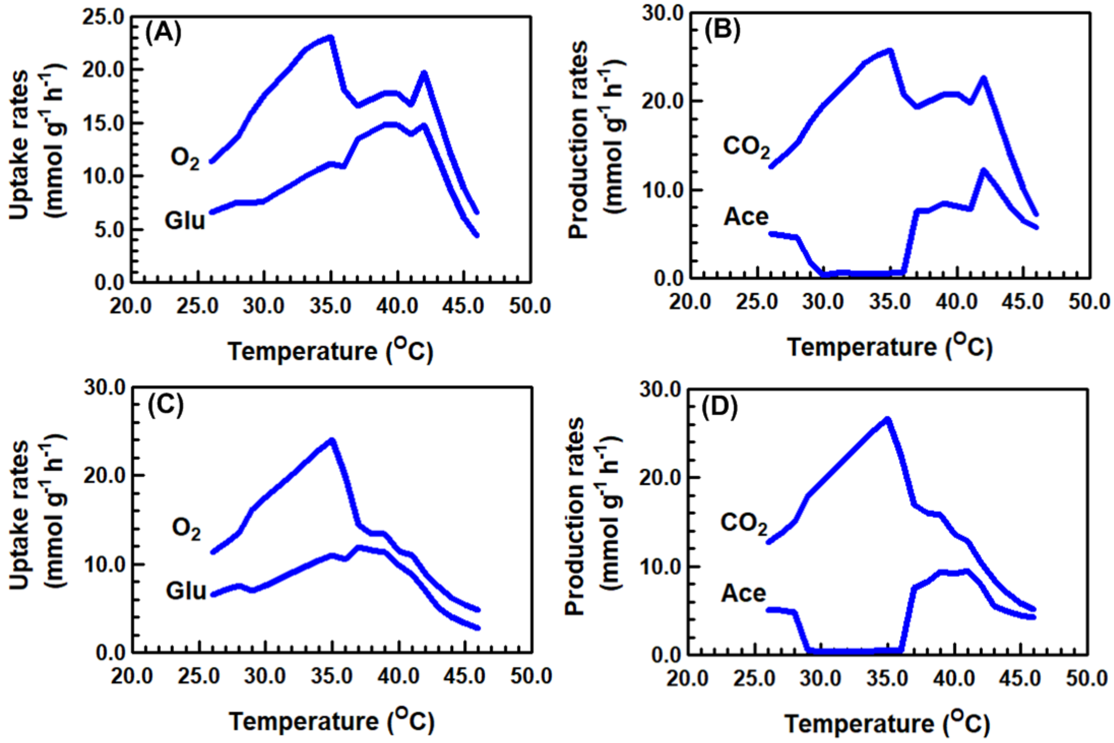


Figure 1S. StressME predicted exchange rates under thermal stress. (A) and (B): heat-evolved strain; (C) and (D): wild-type strain. Exchange rates shown are glucose (Glu) uptake rates, oxygen ($O_2$) uptake rates, carbon dioxide (CO2) production rates (CO2) and acetate (Ace) production rates.

The analysis of the proteome reallocation for the heat-evolved strain at 32 and 40 $^oC$ provides a global overview of how cells adjust their protein synthesis for different metabolic processes when exposed to environmental stress (Figure 2S). As compared to the proteome at 32 $^oC$, more proteome resources at 40 $^oC$ were directed to synthesize protective chaperones (e.g., DnaJ, DnaK, GroS, GrpE) to assist folding and unfolding, sigma factor to regulate the stress response (e.g. rpoN and rpoH), and PPP (pentose phosphate pathway) proteins to maintain carbon homoeostasis (e.g. gnd, rpe and tktB). Meanwhile, cells at 40 $^oC$ invested less proteome resources in energy production (i.e., TCA cycle with succinate dehydrogenase coupled to the electron transport chain) to reallocate the proteome resources into the protective proteins to tolerate the stress conditions.

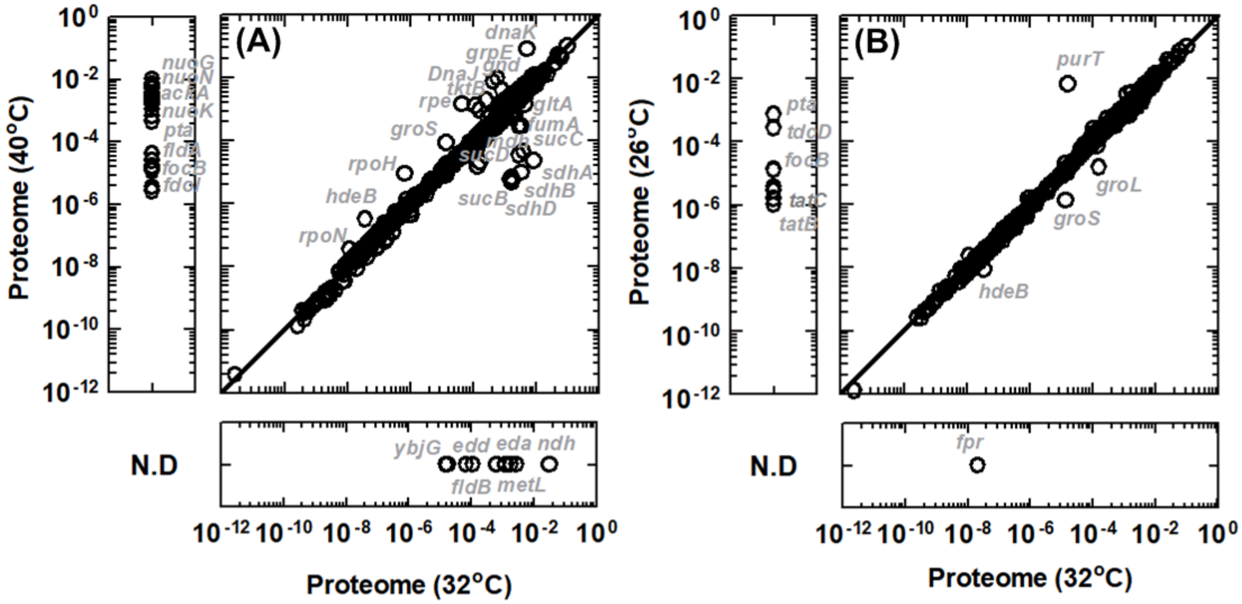


Figure 2S StressME predicted proteome reallocation at different temperatures (A) proteome at 32 ºC and 40 ºC (B) proteome at 32 ºC and 26 ºC

### SI-4: Alternative optima captured by StressME (purT vs. ackA)

**Table S4. Structure of three proteins with any one the three capable of catalyzing ACKr***

| PurT | ackA | tdcD |
|---|---|---|
| ***Number of residues:*** | ***Number of residues:*** | ***Number of residues:*** |
| 392 | 400 | 402 |
| ***Molecular weight (in H2O):*** | ***Molecular weight (in H2O):*** | ***Molecular weight (in H2O):*** |
| natural abundance 42432.84 | natural abundance 43289.60 | natural abundance 43383.28 |
| 2H 44764.45 | 2H 45645.03 | 2H 45690.08 |
| 13C 44289.76 | 13C 45187.08 | 13C 45262.95 |
| 15N 42960.28 | 15N 43811.09 | 15N 43924.63 |
| 2H,13C 46621.37 | 2H,13C 47542.51 | 2H,13C 47569.75 |
| 2H,15N 45291.89 | 2H,15N 46166.51 | 2H,15N 46231.43 |
| 13C,15N 44817.20 | 13C,15N 45708.56 | 13C,15N 45804.30 |
| 2H,13C,15N 47148.81 | 2H,13C,15N 48063.99 | 2H,13C,15N 48111.10 |
| ***Atomic composition:*** | ***Atomic composition:*** | ***Atomic composition:*** |
| Carbon (C) 1877 | Carbon (C) 1918 | Carbon (C) 1900 |
| Hydrogen (H) 3019 | Hydrogen (H) 3051 | Hydrogen (H) 3017 |
| non-exchangeable 2350 | non-exchangeable 2374 | non-exchangeable 2325 |
| exchangeable 669 | exchangeable 677 | exchangeable 692 |
| Nitrogen (N) 531 | Nitrogen (N) 525 | Nitrogen (N) 545 |
| Oxygen (O) 560 | Oxygen (O) 582 | Oxygen (O) 586 |
| Sulfur (S) 14 | Sulfur (S) 16 | Sulfur (S) 16 |

*Protein structure determined based on the amino acid sequence from https://spin.niddk.nih.gov/clore/

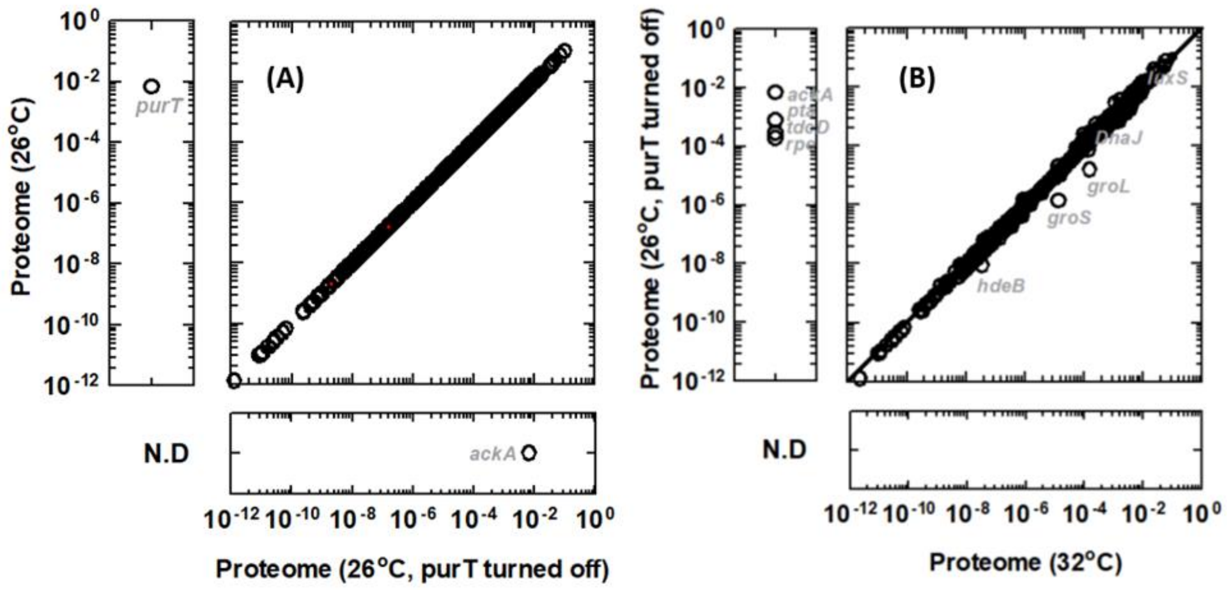


Figure 3S. Effect of purT and ackA translation and protein synthesis on overall proteome.

### SI-5: Reactions for NADH dehydrogenase and Quinolinate synthase

##### NADH dehydrogenase affected by ROS damage

***(a) NADH16pp (NADH dehydrogenase coded by nuo)***

9.27286647516804e-7*mu NADH-DHI-CPLX_mod_2fe2s_mod_4fe4s_mod_fmn + 4.0 h_c + nadh_c + q8_c --> **-3.33823193106049*h2o2_c force_damage_NADH-DHI-CPLX_mod_2fe2s_mod_4fe4s_mod_fmn_h2o2** + **-3338.23193106049*o2s_c force_damage_NADH-DHI-CPLX_mod_2fe2s_mod_4fe4s_mod_fmn_o2s** + 3.0 h_p + nad_c + q8h2_c

***(b) NADH17pp (NADH dehydrogenase coded by nuo)***

9.27286647516804e-7*mu NADH-DHI-CPLX_mod_2fe2s_mod_4fe4s_mod_fmn + 4.0 h_c + mqn8_c + nadh_c --> **-3.33823193106049*h2o2_c force_damage_NADH-DHI-CPLX_mod_2fe2s_mod_4fe4s_mod_fmn_h2o2** + **-3338.23193106049*o2s_c force_damage_NADH-DHI-CPLX_mod_2fe2s_mod_4fe4s_mod_fmn_o2s** + 3.0 h_p + mql8_c + nad_c

##### NADH dehydrogenase unaffected by ROS damage

***(c) NADH5 (NADH dehydrogenase coded by ndh)***

5.65630208035663e-6*mu NADH-DHII-MONOMER_mod_mg2_mod_cu_mod_fad + h_c + nadh_c + q8_c --> nad_c + q8h2_c

##### Quinolinate synthase affected by ROS damage

***(d) QULNS (quinolinate synthase coded by nadA)***

0.01984448692471*mu CPLX0-7719_mod_4fe4s + dhap_c + iasp_c --> **-71440.1529289559*h2o2_c force_damage_CPLX0-7719_mod_4fe4s_h2o2** + **-71440152.9289559*o2s_c force_damage_CPLX0-7719_mod_4fe4s_o2s** + 2.0 h2o_c + pi_c + quln_c

### SI-6: StressME using Linux (clusters)

#### Installation

```
# To set up virtual environment for python 3.6


virtualenv -p python MePython363
source MePython363/bin/activate

# install dependencies
```

| Accessories | Version | Installation |
|---|---|---|
| (a) cython | 0.28.2 | pip |
| (b) sympy | 1.1.1 | pip |
| (c) numpy | 1.14.3 | pip |
| (d) scipy | 1.1.0 | pip |
| (e) pytest | 3.5.1 | pip |

| | | |
|---|---|---|
| (f) pandas | 0.22.0 | pip |
| (g) cycler | 0.11.0 | pip |
| (h) matplotlib | 2.2.2 | pip |
| (i) biopython | 1.76 | pip |
| (j) qMINOS* | 5.6 | https://github.com/SBRG/solvemepy |
| (k) cobrame | StressME 1.1 | https://github.com/QCSB/StressME |
| [l] ecolime | StressME 1.1 | https://github.com/QCSB/StressME |
| [m] oxidizeme | StressME 1.1 | https://github.com/QCSB/StressME |
| [n] acidifyme | StressME 1.1 | https://github.com/QCSB/StressME |
| [o] meuser | StressME 1.1 | https://github.com/QCSB/StressME |

*See https://github.com/SBRG/solvemepy for qMINOS installation

**Simulations on Linux clusters by slurm**

*salloc --time=2:0:0 --ntasks=1 --cpus-per-task=1 --mem-per-cpu=8G --account=<your_account> python StressME_wildtype.py 42 5.0 10*

or:

*sbatch --mem=8G --account=<your_account> --time=2:00:00 --output StressME_wildtype StressME_wildtype.sh*

where StressME_wildtype.sh is coded as:

#!/bin/bash

#SBATCH --time=2:00:00
#SBATCH --account=<your_account>
python StressME_wildtype.py 42 5.0 10

Here "42 5.0 10" refers to the triple stress conditions at temperature 42 °C, pH 5.0 and ROS 10X of the basal level.

### SI-7: StressME using docker

Docker allows ME-model users to run StressME locally without going through the complicated processes of installing solvers and dependencies (SI-6) that may be incompatible with each other due to their version update.

The Docker image of StressME can be found on Docker Hub (queensysbio/stressme:v1.1). This build was developed from the modified version of COBRAme and EcoliME kernel to integrate FoldME and AcidifyME with OxidizeME. This build also includes qMINOS solver that users can use to solve StressME using solvemepy.

The installed COBRAme (StressME version), ECOLIme (StressME version), AcidifyME, OxidizeME and solvemepy packages can be found in /source/ after a new container has been created from the image of StressME. The working directory is /home/meuser, where ME-model users can run simulations and export output from the StressME container to the host.

##### Installation on Windows Subsystem for Linux (WSL2)

Steps to run a Docker container from the image of StressME with everything required to run the StressME using qMINOS:

(a) Install Ubuntu on WSL2 on Windows 10/11. https://ubuntu.com/tutorials/install-ubuntu-on-wsl2-on-windows-10#1-overview
(b) Download Docker Desktop for Windows https://desktop.docker.com/win/main/amd64/Docker%20Desktop%20Installer.exe
(c) Start Docker Desktop from the Windows Start menu. From the Docker menu, select **Settings** and then **General**. Select the **Use WSL 2 based engine** check box. Select **Apply & Restart**
(d) Pull the latest StressME image from DockerHub with docker pull queensysbio/stressme:v1.1.

##### Simulations on Windows Subsystem for Linux (WSL2)

(a) Start Docker Desktop from the Windows Start menu
(b) Start another PowerShell window from the Windows Start menu. In the command line, type "ubuntu"
(c) Type "cd /mnt/c/Users/<your user name in this local computer>/Desktop" and then type "mkdir StressME_wild_simulations" and "mkdir StressME_heat_simulations" to create host directories to receive output from the Docker StressME container
(d) Type "cd StressME_wild_simulations" or "cd StressME_heat_simulations" and then type "docker run -p 8888:8888 --rm -i -v $(pwd):/mount_point/ -t queensysbio/stressme:v1.1 bash" . This will start the Docker StressME container (virtual machine) in the working directory: /home/meuser.
(e) Type "ls" to check files available in the working directory.
(f) Type "python StressME_wildtype.py 42 5.0 10" or "python StressME_heatevolved.py 42 5.0 10" to run simulations for the wild type or the heat evolved strains. Here "42 5.0 10" refers to the triple stress conditions at temperature 42 °C, pH 5.0 and ROS 10X of the basal level.
(g) After simulations are done, three csv files (TripleStressME_proteome.csv, TripleStressME_phenotypes.csv, and TripleStressME_fluxes.csv) are found in the working directory /home/meuser, representing the protein mass fractions, metabolic fluxes and phenotypes under the test stress conditions. Type "cp *.csv /mount_point/ StressME_wild_simulations" or "cp *.csv /mount_point/ StressME_heat_simulations" to export results to the corresponding host directories.
(h) Use "CTRL+D" to exit Docker StressME container to go back to the host directory.
(i) Type "explorer.exe ." to open File Explorer to view (e.g. EXCEL) and process (e.g., R for Rstudio or Matlab) the csv data.

##### Simulations on Jupiter notebook

(a) Start Docker Desktop from the Windows Start menu
(b) In WSL2 or powershell, run ‘jupyter notebook --ip=0.0.0.0 --port=8888 --allow-root’
(c) Copy the random token that is generated on the screen.
(d) Paste the token on http://localhost:8888/ to launch jupyter notebook
(e) Run StressME_heatevolved.ipynb and StressME_wildtype.ipynb for simulations.
